# Transcriptomic analysis of four cerianthid (Cnidaria, Ceriantharia) venoms

**DOI:** 10.1101/2020.06.18.159541

**Authors:** Anna M. L. Klompen, Jason Macrander, Adam M. Reitzel, Sérgio N. Stampar

## Abstract

Tube anemones, or cerianthids, are a phylogenetically informative group of cnidarians with complex life histories, including a pelagic larval stage and tube-dwelling adult stage, both known to utilize venom in stinging-cell rich tentacles. Cnidarians are an entirely venomous group that utilize their proteinaceous-dominated toxins to capture prey and defend against predators, in addition to several other ecological functions, including intraspecific interactions. At present there are no studies describing the venom for any species within cerianthids. Given their unique development, ecology, and distinct phylogenetic-placement within Cnidaria, our objective is to evaluate the venom-like gene diversity of four species of cerianthids from newly collected transcriptomic data. We identified 525 venom-like genes between all four species. The venom-gene profile for each species was dominated by enzymatic protein and peptide families, which is consistent with previous findings in other cnidarian venoms. However, we found few toxins that are typical of sea anemones and corals, and furthermore, three of the four species express toxin-like genes closely related to potent pore-forming toxins in box jellyfish. Our study is the first to provide a survey of the putative venom composition of cerianthids, and contributes to our general understanding of the diversity of cnidarian toxins.

## 1. Introduction

The phylum Cnidaria (sea anemones, corals, jellyfish, box jellies, hydroids/hydromedusae, etc.) is the earliest diverging venomous lineage (∼ 600 million years) [1,2]. Cnidaria deliver their proteinaceous-dominant venom through organelles called nematocysts (a type of cnidae), housed in cells called nematocytes [3,4]. Venom from discharged nematocysts is used in prey capture and defense against predation, but cnidarians also use venom for a variety of other behaviors, such as intraspecific competition [5-7] and maternal care [8] (see review by [9]). This ecological diversity is complemented by the functional diversity of cnidarian venoms, which can include neurotoxic, cytotoxic, and enzymatic (e.g. phospholipase and metalloprotease) proteins and peptides, in addition to non-peptidic components [10,11]. For humans, stings from certain species can cause intense localized pain, scarring, induced anaphylaxis, and in the worst cases, cardiac and respiratory failure leading to death [12-15]. The venom of medically relevant species, such as the Portuguese Man-o-War (*Physalia physalis*) [16-18] and several species of box jellyfish ([19-22], reviewed in [23]), or easy to collect species, such as sea anemones [24,25], have been explored more extensively at a biochemical and pharmacological level [26]. However, these species represent a small fraction of the species diversity within the group, and only recently has the exploration the venom composition for a wider number of cnidarians increased in an effort to characterize the evolution and ecological function of toxins within the group [27].

There is also a growing interest in cnidarian venoms as a potential resource for drug discovery, particularly the neurotoxin-rich venoms of sea anemones [28-30]. One of the best studied therapeutic proteins derived from a cnidarian toxin is an analogue of a potassium Kv1.3 channel blocker isolated from the sun sea anemone (*Stichodactyla helianthus*) called ShK [31], which completed Phase 1b trials for autoimmune diseases [32,33]. Because ShK-scaffolds are abundant in sea anemone venom peptides, characterizing the venoms from sea anemones (and cnidarians in general) could yield additional candidates for novel therapeutic compounds [30,34,35]. Kunitz-domain containing serine inhibitors, also found in sea anemone venoms, can also be used as potential therapeutic resources [25,36]. These cnidarian-derived neuropeptide inhibitors have potential applications as analgesics, antiepileptics, and other neuroprotective drugs [37].

While there has been a recent increase in transcriptomic and proteotranscriptomic analyses of cnidarian venoms (e.g. [7,8,22,38-54]), the phylum as a whole, which contains over 13,000 species, remains highly understudied. Cnidaria is split into three taxonomic groups: Anthozoa (sea anemones, corals, zoanthids, etc.), Medusozoa (jellyfish, box jellies, hydroids, siphonophores), and Endocnidozoa (*Polypodium* + myxozoans) [55,56]. Of the 7,142 animal toxins and venoms listed in Tox-Prot, a curated animal venom annotation database, only 273 are derived from cnidarians (as of May 2020, [57]), with that vast majority (>96%) are from anthozoans. Within that limited number there is even greater taxonomic bias; almost 90% of anthozoan toxins are from the Actinioidea superfamily of sea anemones [27,30], meaning less than 50 out of 1,100 known sea anemone species contribute to the database of annotated cnidarian toxins [54]. This taxon bias limits researcher’s ability to discover novel therapeutic peptides and scaffolds from sea anemones, as well as limits to search for potential drug candidates in other anthozoan groups such as corals [58] and zoanthids [47-49].

One major hurdle to identifying the composition and comparative diversity of cnidarian toxins is their lack of a centralized venom system that can be easily isolated for study. This packaging of toxins into individual nematocysts scattered throughout the animal, impedes the ability to isolate crude venoms for downstream analysis, which is further exacerbated in smaller or rare species of cnidarians. There are several protocols for isolating venom from nematocysts (e.g. [59-62]), but these methods, as noted above, are typically restricted to larger or easy to obtain animals (e.g. corals and sea anemones, true jellies such as *Chrysaora* and *Cyanea*), species of medical relevance (e.g. *Physalia*, box jellies), or those that can be easily maintained in a lab (e.g. *Hydra* [63], *Nematostella* [64]). Next generation sequencing technologies provide a solution to this problem, and have greatly increased the ability of researchers to screen the diversity of putative venom-like genes for neglected or poorly studied venomous species, including cnidarians [65].

One group of anthozoans whose venoms have yet to be explored are members of the subclass Ceriantharia, known as cerianthids (Phylum Cnidaria: Class Anthozoa) (Figure 1). Cerianthids are tube-dwelling anemones, so named because of their ability to create complex tubing from a specialized group of cnidae called ptychocysts [66]. Their phylogenetic placement within Cnidaria remains contentious, due to a combination of a lack of available sequence data and low species sampling [5,67,68]. Various studies place them as sister group to Hexacorallia, sister group to Octocorallia [69], or sister group to Hexacorallia + Octocorallia [70,71]. Although cerianthids are clearly members of Anthozoa, they have several features that are more similar to Medusozoa. For instance, cerianthids possess linear mitochondrial genomes, as in medusozoans, while all other anthozoans have circular mitochondrial genomes [71-73]. Also, unlike other anthozoans, cerianthids display a long-lived pelagic larval stage that superficially resembles a medusa [74]. It is unclear how this unique life history or their early diverging phylogenetic relationship to either, or both, of the major groups of anthozoans may be reflected in the venom composition of this group relative to other anthozoan venoms (or cnidarians more generally).

**Figure 1.**
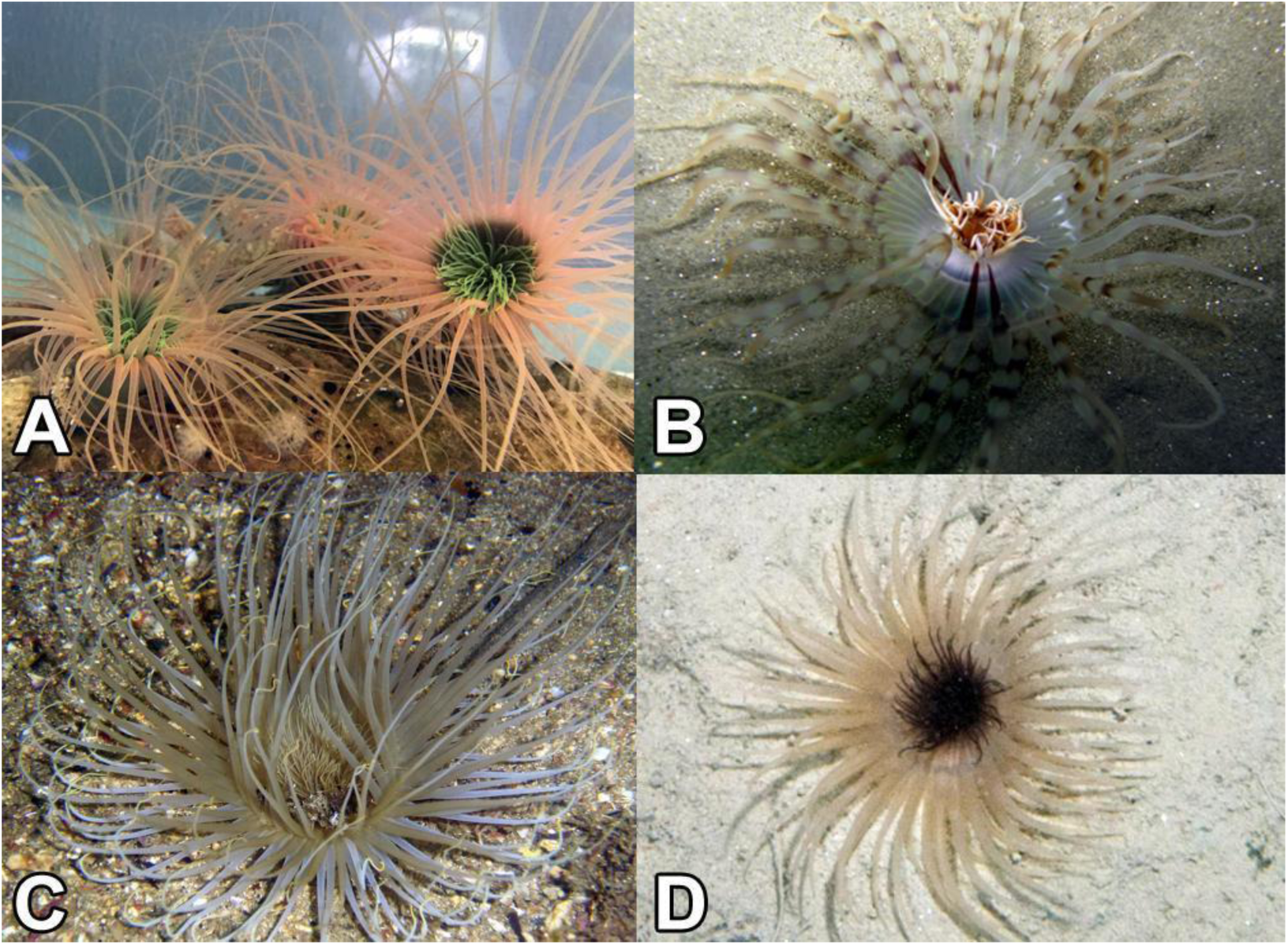
Ceriantharia species used in the current study. A) *Pachycerianthus cf. maua*; B) *Isarachnanthus nocturnus*; C) *Ceriantheomorphe brasiliensis* and D) *Pachycerianthus borealis*. Photos by Fisheries and Oceans Canada (Claude Nozères)).

The aim of this project is to explore newly sequenced transcriptomes for four adult cerianthid species (*Ceriantheomorphe brasiliensis, Isarachnanthus nocturnus, Pachycerianthus borealis*, and *Pachycerianthus cf. maua*) and determine putative venom-like gene candidates across each using a customized annotation pipeline. This study is the first formal analysis of venom composition within this subclass Ceriantharia, and a targeted comparison of the venom gene profiles between cerianthids and other cnidarian species.

## 2. Results

### 2.1. Results for sequencing and de-novo transcriptome assembly of four cerianthids species

The number of paired end reads generated by Illumina HiSeq runranged from 27,865,720 to 36,520,791 across all taxa. The Trinity [75] assembly ranged from 92,757 to 158,663 unique assembled transcripts with an N50 range from 1101 - 1282. Overall completeness evaluated in BUSCO ranged from 88.1% to 97.9% complete (Table 1).

**Table 1.** Sequencing and assembly parameters for various cerianthid transcriptomes.

| Species | Reads (PE) | Transcripts | Genes | N50 | BUSCO % |
| --- | --- | --- | --- | --- | --- |
| *C. brasiliensis* | 34,877,883 | 131,550 | 110,524 | 1,276 | 95.4% |
| *I. nocturnus* | 31,028,274 | 92,757 | 78,821 | 1170 | 89.2% |
| *P. borealis* | 36,520,791 | 158,633 | 120,542 | 1,282 | 97.9% |
| *P. maua* | 27,865,720 | 179,576 | 145,788 | 1101 | 88.1% |

### 2.2. Diversity and phylogenetic analysis of putative venom-like gene profiles for cerianthids species

Using the de-novo assemblies, we identified a diverse set of venom-like putative protein coding transcripts and peptides across the four cerianthids: 169, 69, 182, and 105 for *C. brasiliensis, I. nocturnus, P. borealis*, and *P. maua*, respectively. All toxins were categorized into families/scaffolds based on their highest Tox-Prot (i.e. UniProtKB/Swiss-Prot) BLAST hit [57], and categorized by biological function: neurotoxin, hemostatic and hemorrhagic toxins, membrane-active toxins, mixed function enzymes, protease inhibitors, allergen and innate immunity, and venom auxiliary proteins (modified from [49]). A summary of annotated contigs for each species is shown in Figure 2, Table Below we provide short descriptions of select toxin groups and families represented by the identified toxins.

**Figure 2.**
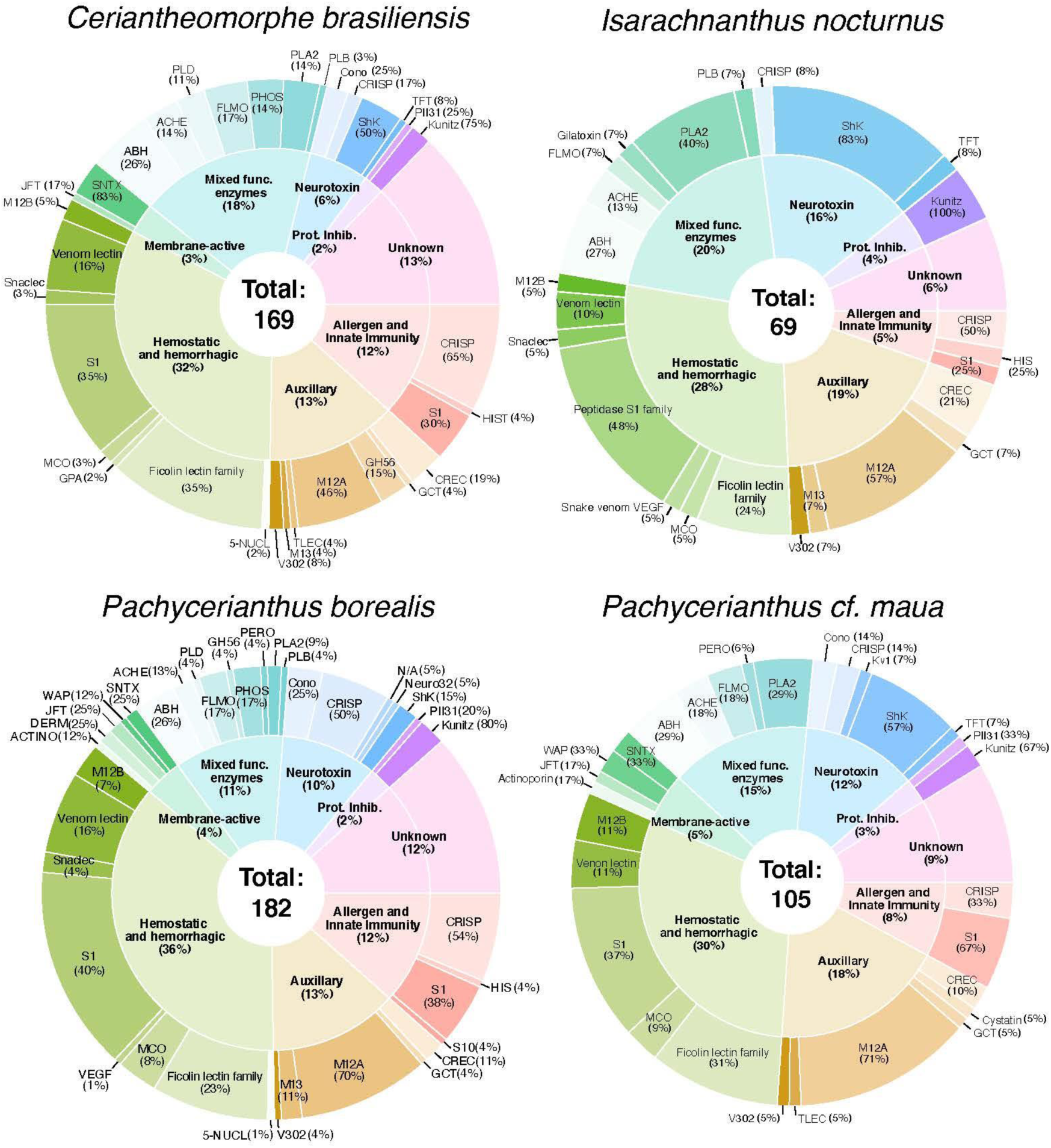
Number of venom-like genes identified for four cerianthid species. Inner circle: biological function and overall percentage of each over the total venom-like gene profile in each species. Outer circle: Venom-like genes families within each biological function category and overall percentage of that family within each category. ABH = AB hydrolase superfamily; ACHE = Acetylcholinesterase; ACTINO = Actinoporin-like; Cono = Conopeptide P–like superfamily; DERM = Dermatopontin; FLMO = lavin monoamine oxidase; GCT = Glutaminyl–peptide cyclotransferase; GH56 = Glycoside hydrolase 56; GPA = Glycoprotein hormones subunit alpha; HIS = Histidine acid phosphatase; JFT = Jellyfish Toxin; Kunitz = Venom Kunitz–type; Kv1 = Sea anemone type 1 potassium channel toxin ; M12A = Peptidase M12; MCO= Multicopper oxidase; M12B = Venom metalloproteinase (M12B); M13 = Peptidase M13; Neuro32 = Neurotoxin 32 Family; PII31 = Protease inhibitor I31; PHOS = Nucleotide pyrophosphatase/phosphodiesterase; PLA2 = Phospholipase A2; PLB = Phospholipase B-like; PLD = Arthropod phospholipase D; PERO = Peroxiredoxin ; SNTX = SNTX/VTX toxin; S1,S10 = Peptidase S1,S10; Venom Lectin = True venom lectin ; TLEC = Techylectin–like; TFT = Snake three–finger toxin; VEGF = Venom vascular endothelial growth factor; V302 = Venom protein 302; WAP = Snake waprin; 5-NUCL = 5’–nucleotidase.

#### 2.1.1. Neurotoxins

ShK-domain containing proteins and peptides are some of the most diverse toxins within the transcriptomes of the four species, which includes 15 cysteine-rich venom proteins, 27 ShK-domain containing toxins as identified from Pfam [76,77] (Supplemental Figures S1, S2), and a single sea anemone type 1 potassium channel toxin in *P. maua*. Interestingly, a single transcript in *P. borealis* that contains an ShK-domain had the closest match to propeptide 332-1 toxin from *Malo kingi*, a box jellyfish with a potent sting known to cause Irukandji syndrome [78]. Though the functions are highly variable and depend on the combination of present domains [30,79], ShK-domain toxins can cause paralysis due to potassium channel inhibition as well as induce hemolytic effects [80,81]. As noted above, these ShK toxins may also confer structural and/or functional properties of interest for pharmacological research.

Turripeptides are ion channel blockers described from turrid gastropods, relatives of cone snails, but they have also been predicted or isolated from three species of zoanthid [47-49], a box jellyfish [22], a true jellyfish [82], and a stalked jellyfish [83], as well as bloodworms and marine annelids [81]. These toxin peptides contain a kazal domain with a conserved cysteine framework (C-C-C-C-C-C), and modulate ion channels, resulting in paralysis [84,85]. Four transcripts from cerianthids were shown to have similar cysteine patterns architecture, but have longer predicted protein sequences than the typical turripeptide sequences of <100 amino acids and four additional conserved cysteines upstream from the kazal domain (Figure 3).

**Figure 3.**
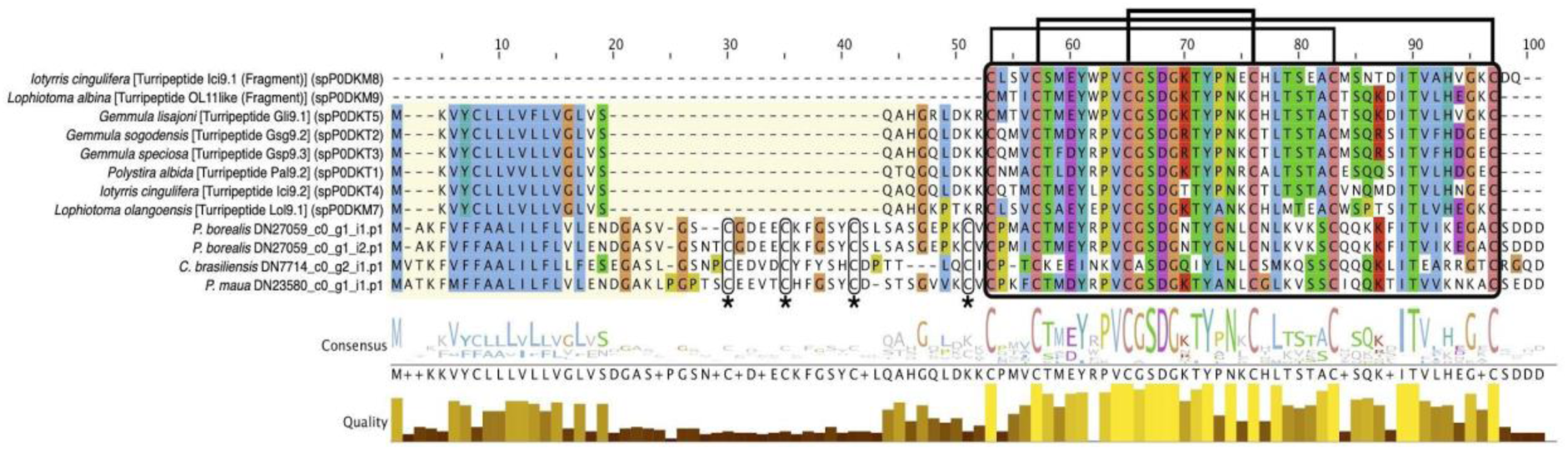
Multiple sequence alignment of candidate turripeptide-like sequences for cerianthid toxins and representatives from cone snails created using L-INS-I algorithm via MAFFT [153], viewed using Jalview [155] with Clustal color scheme. Kazal domain (in black box) and conserved cysteine patterning shown (bridging) are highlighted. The yellow box indicates the predicted signal peptide sequences as indicated by Signalp [147]. The stars and corresponding smaller black boxes indicate the four cysteine residues that are present in the cerianthid sequences preceding the kazal domain.

Three sequences, one each from *C. brasiliensis, I. nocturnus*, and *P. maua*, closely matched to three-finger toxins (TFTs), snake-derived toxins that display a wide diversity of functions such as neurotoxicity, acetylcholinesterase inhibitors, cytotoxins (cardiotoxins), platelet aggregation inhibitors, coagulation factor inhibitors, heparin binders, and K+ channel, and integral-receptor ligands [86]. A recent proteomic study found that the orange cup coral *Tubastrea coccinea* contains a putative TFT toxin [83], in addition to a predicted TFT in *P. varibilis* [47]. The TFT toxins in cerianthids and *P. varibilis* cluster as sister to bucandin, a TFT isolated from Malayan krait (*Bungarus candidus*) [87]. However, the bootstrap support throughout the phylogeny is generally low (<70%) (Supplemental Figures S4).

#### 2.2.2. Hemostatic and hemorrhagic toxins

Hemostatic and hemorrhagic toxins are the most diverse type of toxins in all four cerianthid species (Figure 2). They generally interfere with hemostasis through various pathways, either individually or synergistically with other toxins. This group includes a variety of C-type lectin-containing toxins (C-type lectin lectoxin, galactose specific lectin, and snake c-type lectin (snaclec)), and are associated with blood coagulation, inflammation, myotoxicity, and homeostasis interference [87,88]. They have been reported in a variety of animal venoms, including, crustaceans, blood feeding insects, caterpillars, leeches, bloodworms, snakes, and stonefish [88], as well as cnidarian species [38,43,44,47,49]. We found 34 total toxins between the four species that match to a C-type lectin domain.

One of the most numerous groups of venom-like genes within this class are putative veficolin-like toxins (total 30), which are, comparatively, highly abundant in *P. borealis* (9 sequences) and *C. brasiliensis* (14 sequences). This toxin was described from the Komodo dragon (*Varanus komodoensis*), and is suggested to interfere with blood coagulation and/or platelet aggregation based on the similarity to ryncolin toxins [90]. Ryncolin toxins are represented in all cerianthid assemblies in relatively high abundance with 25 total sequences, originally described from the dog-faced water snake (*Cerberus rynchops*). Six sequences from the transcriptome of the zoanthid *Palythoa caribaeorum* (categorized in our study under allergen and innate immunity) [48] and three peptides from the proteome of the scyphozoan *Nemopilema nomurai* (as *Stomolophus meleagris*) [38] also belong in this group, suggesting ryncolin-like toxins may play be present across cnidarians.

We also found numerous venom prothrombin activators in two different groups: Factor 5/8 C-domain and trypsin domain. These types of toxins are well known from snake venoms, and cause hemostatic impairment by proteolytic cleavage of prothrombin to thrombin [91]. Putative transcripts have been found in relatively high abundance in the mat anemone *Zoanthus natalensis* [49] as well as in the transcriptomes of *P. caribaeorum* [48] and sea anemone *Anthopleura dowii* [53]. They have also been found in a transcriptomic analysis of the sea anemone *Stichodactyla haddoni* venom, but no peptides were detected using mass spectrometry [46], suggesting that additional proteomic experiments will be needed to confirm the presence of these prothrombin activators (and other toxin groups) in cerianthid venoms.

#### 2.2.3. Membrane-active toxins, protease inhibitors

Jellyfish toxins (or CaTX/CrTX) are one of the most potent toxin families from cnidarians, initially isolated from several species of box jellyfish possessing stings that are dangerous to humans [20]. Two members within this family, CfTX-1 and CfTX-2 from the Australian box jellyfish (*Chironex fleckeri*), are highly cardiotoxic, and their stings are associated with cardiac failure [41]. Four sequence from cerianthids, two from *P. borealis* and one each from *C. brasiliensis* and *P. maua*, appear to belong in this family based on strong phylogenetic evidence, although the transcript from *P. maua* clustered with toxins from the hydroid *Hydra vulgaris* [92], which have yet to be functionally analyzed (Figure 4).

**Figure 4.**
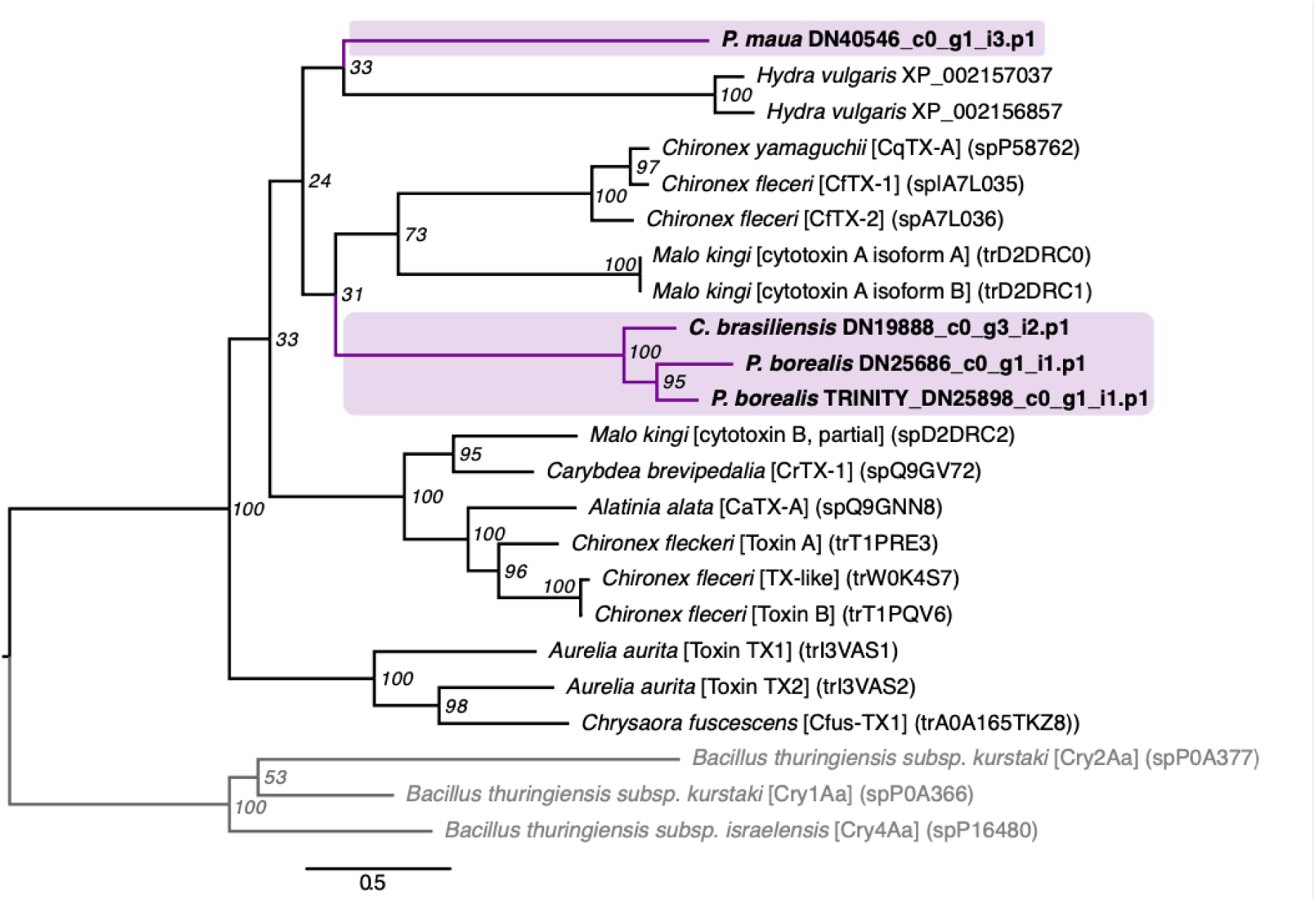
Phylogenetic tree of jellyfish toxin (or CaTX/CrTX) sequences. Phylogeny was constructed using RAxML with the PROTGAMMAWAG option [154]. Bootstrap support based on 500 rapid bootstrap replicates, and all support values are shown. Putative genes outlined in purple are from cerianthids sequences. Sequences in gray are bacterial pore-forming toxins that have closest structural homology to this toxin family [14], used to root the tree.

Originally derived from sea anemones, actinoporins are conserved 20kDa pore-forming toxins that exhibit cytolytic and hemolytic effects [93]. Actinoporin-like sequences have also been isolated from both molluscs [94] and chordates [95], and shown to be toxic to a wide variety of vertebrate and invertebrate species [96,97]. Two actinoporin sequences similar to DELTA-thalatoxin-Avl2a were found in *P. borealis* and *P. maua*, though both were phylogenetically closer to actinoporin-like sequences found in venomous gastropods and a putative actinoporin from *P. varibilis* [47] (Figure 5).

**Figure 5.**
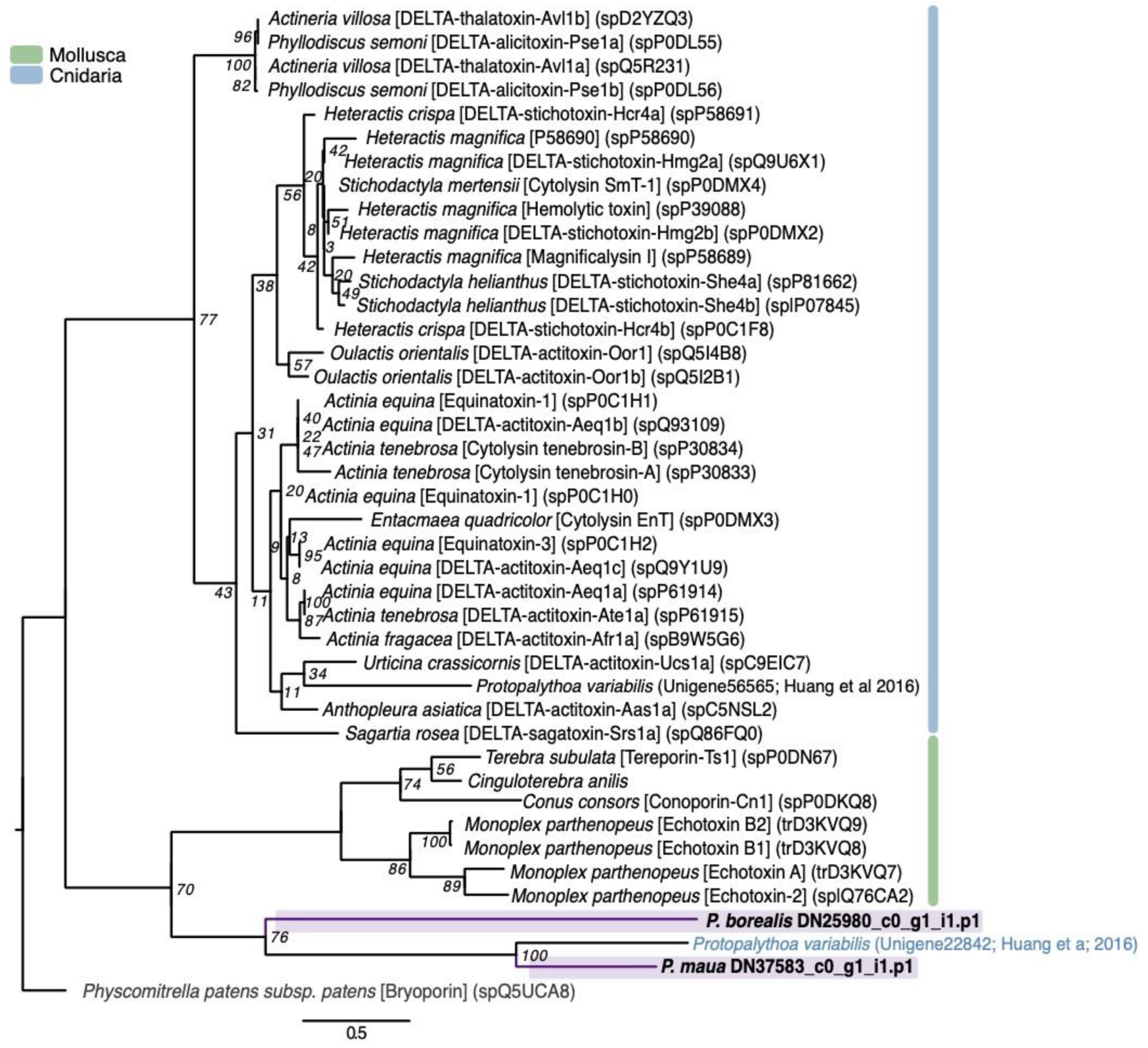
Phylogenetic tree of actinoporin and actinoporin-like sequences. Phylogeny was constructed using RAxML with the PROTGAMMAWAG option [154]. Bootstrap support based on 500 rapid bootstrap replicates, and all support values are shown. Putative genes outlined in purple are from cerianthids sequences. Sequences in gray are non-venomous representatives, and other colors outlined in the key are venom-like genes from other animal classes. Phylogeny modified from von [81]. Tree is rooted with actinoporin-like sequence from a moss (*Physcomitrella patens subsp. patens*).

SNTX-like transcripts include stonutoxin and neoverrucotoxin, non-enzymatic proteins found in a diversity of scorpaeniform fish and monotremes mammals [98,99]. In fish, these toxins cause lethal hemolysis and disrupt circulatory and neuromuscular systems [100,101]. *P. borealis, C. brasiliensis*, and *P. maua* express 9 SNTX-like transcripts, all of which phylogenetically cluster together in a group with two SNTX-like genes from non-venomous fish that is sister to a clade of SNTX genes from highly toxic stonefish (Figure 6).

**Figure 6.**
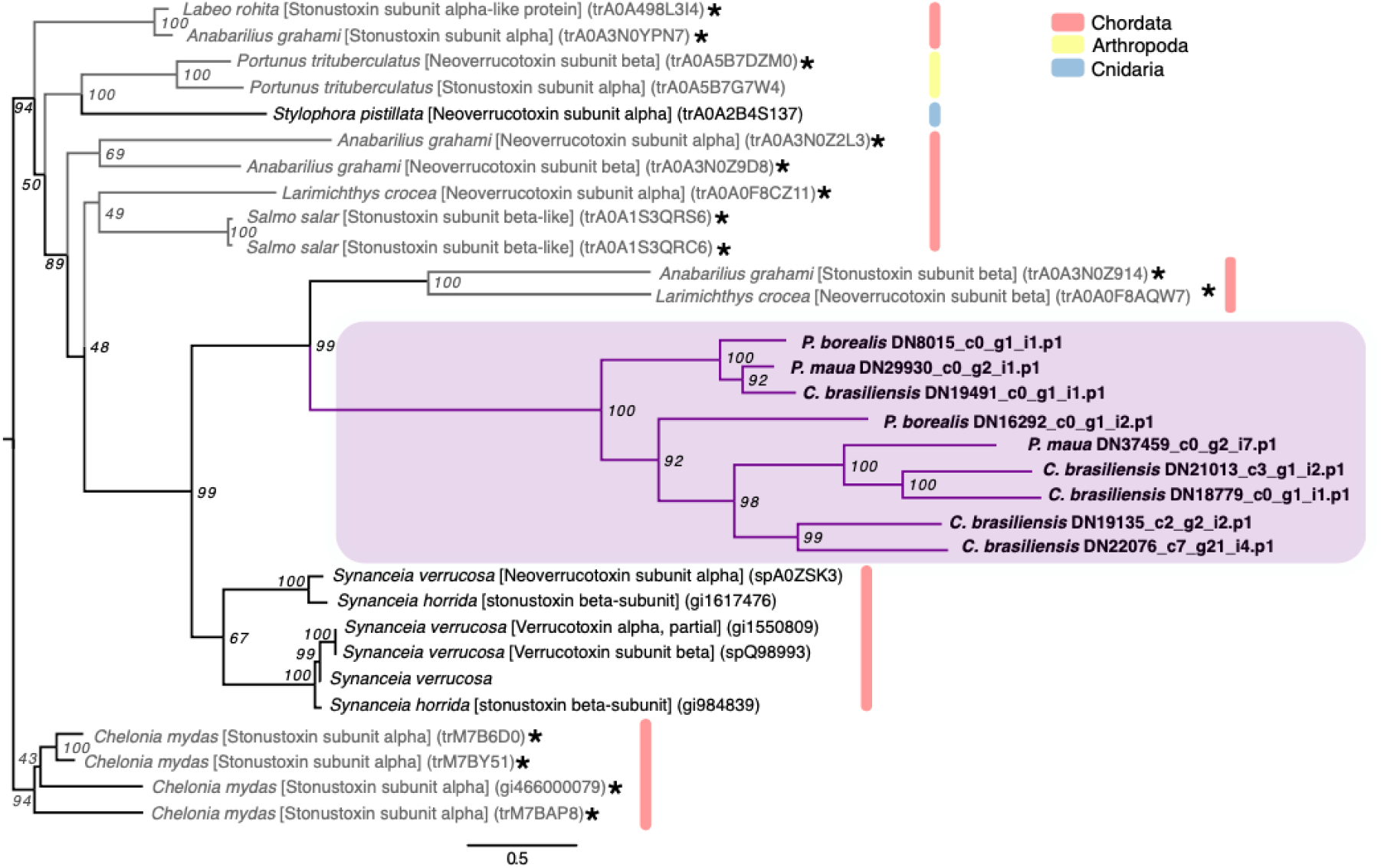
Phylogenetic tree of SNTX-like family sequences. Phylogeny was constructed using RAxML with the PROTGAMMAWAG option [154]. Bootstrap support based on 500 rapid bootstrap replicates, and all support values are shown. Putative genes outlined in purple are from cerianthids sequences. Sequences in gray and starred are non-venomous representatives, and other colors are from other animal classes. Phylogeny modified from [81]. Tree is rooted with sequences from green sea turtle (*Chelonia mydas*).

Waprins are membrane-active toxins derived from snakes that act as antimicrobial proteins, which are used by venomous animals as a defense against microbial infections of their venom glands [102,103]. One sequence of a waprin-like toxin from *P. borealis* and two from *P. maua* were identified in the cerianthids.

#### 2.2.4. Mixed function enzymes

Phospholipases hydrolyze phospholipids to fatty acids and lysophospholipids, which in venoms induced hemolysis [104,105], as well as tissue necrosis, inflammation, blood coagulation inhibition, and neuromuscular transmission blockage [88,105]. These lipases are found in many animal venoms, including cephalopods, insects, spiders, scorpions, and reptiles [88]. Phospholipase A2 (PLA2) is a common and often abundant enzyme in cnidarians venom that aids in prey capture and digestion, and appears to have antimicrobial activity [106]. PLA2 are the most diverse of the enzymatic toxins detected in cerianthids, with 18 total sequences. Of these, 16 phylogenetically form a cluster that includes a putative PLA2 from *P. variabilis* [47] and conodipine-M alpha chain toxin, which was derived from the Magician’s cone snail (*Conus magus*) and inhibits the binding of isradipine to L-type calcium channels [107] (Figure 7). The other two genes from *C. brasiliensis* and *I. nocturnus* cluster with a PLA2 from the broadclub cuttlefish (*Sepia latimanus*). We additionally found three phospholipase-B toxins within *P. borealis, C. brasiliensis*, and *I. nocturnus* and five phospholipase-D toxins, four in *C. brasiliensis* and a single transcript in *P. borealis*. Phospholipase-D in particular is thought to contribute to the dermonecrotic effects of brown spider venoms [108].

**Figure 7.**
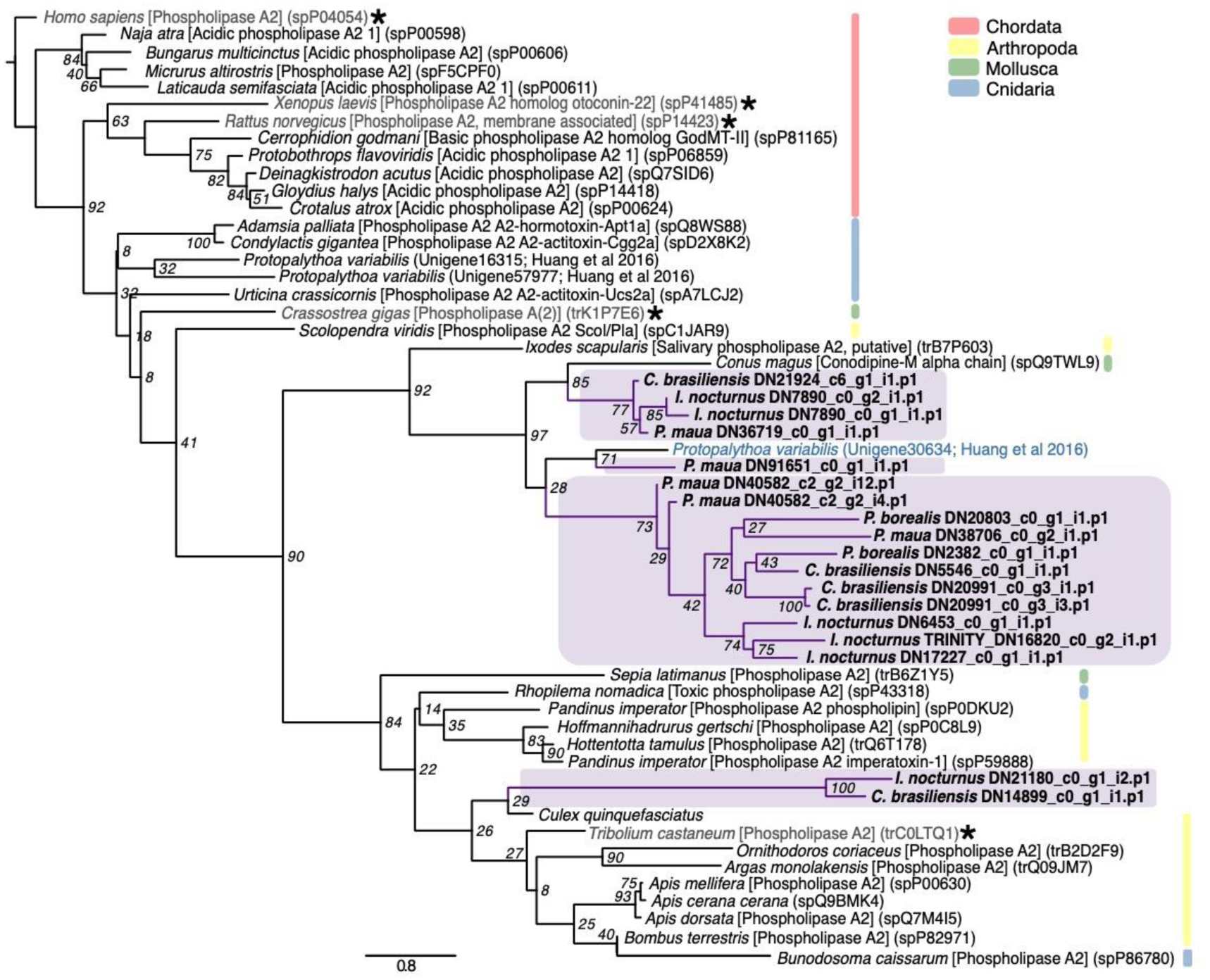
Phylogenetic tree of phospholipase A2 family sequences. Phylogeny was constructed using RAxML with the PROTGAMMAWAG option [154]. Bootstrap support based on 500 rapid bootstrap replicates, and all support values are shown. Putative genes outlined in purple are from cerianthids sequences. Sequences in gray and starred are non-venomous representatives, and other colors are from other animal classes. Phylogeny modified from [81].

#### 2.2.5. Protease inhibitors

Kunitz-domain peptides both block ion channels and inhibit proteases, which can cause blood coagulation, fibrinolysis, and inflammation [109]. In sea anemones, kunitz-containing peptides are typically classified as type II potassium channel toxins, which cause paralysis by blocking potassium channels [25]. All four species have at least one kunitz-type serine protease inhibitor (total 11 across all four species), and *P. maua* specifically has a transcript that matches the sea anemone specific kunitz-containing toxin U-actitoxin-Avd3m, which, based on sequence similarity to other known toxins, may display hemolytic activity as well as potassium channel inhibition.

Three cerianthids, *P. borealis, C. brasiliensis*, and *P. maua* each contain a single transcript that corresponds to a ctenitoxin. Ctenotoxins are thyroglobulin type-1 protease inhibitors originally derived from the Brazilian spider (*Phoneutria nigriventer*), which inhibits cysteine proteases, aspartic proteases and metalloproteases [110].

#### 2.2.6. Allergen and innate immunity

Several components from cnidarian stings have been known to cause immunological responses [14,111]. One common domain of these toxins is the CAP domain, which includes cysteine-rich secretory proteins (CRISPs), antigen 5 (Ag5), and pathogenesis-related 1 (Pr-1) proteins [112]. These are found in many venomous taxa such as cephalopods, bloodworms, fireworms, scorpions, spiders, and reptiles [81,88,113], and are commonly found in cnidarians [22,43]. Function appears to vary by taxonomic group; in snakes, CAP proteins act as ion channel blockers and inhibit smooth muscle contraction [114], in cone snails as proteolytic compounds [115], and in hymenopterans as allergens [116]. The majority of CAP-domain cerianthid transcripts belong to a group called venom allergen proteins (total 31), though this is mainly driven by the number of genes present in *P. borealis* (12 sequences) and *C. brasiliensis* (14 sequences). Both species also have an additional CAP-domain (CRISP/Allergen/Pr-1) toxin. Multiple venom allergen proteins were also reported in the venom of the Pacific sea nettle (*Chrysaora fuscescens*) [43].

#### 2.2.7. Venom auxiliary proteins

Venom auxiliary proteins are secreted in the venom gland to facilitate proper processing and stabilization. They can also work synergistically with other venom components to facilitate the spread of toxins after envenomation. One example is venom protein 302, originally derived from the scorpion *Lychas mucronatus* [117]. Each cerianthid has a putative single venom protein 302 match, two in the case of *C. brasiliensis*, but (weak) phylogenetic signals suggests that the cerianthids proteins are more closely related to an insulin-like growth factor-binding (IGLFP) protein from hexacorallian *S. pistillata* [118] (Supplementary Figure S9). Two venom protein 302 proteins were also identified in *P. variabilis* [47], and these zoanthid toxins formed a clade that is a sister group to non-venomous IGLFP-domain containing proteins in our study (Supplementary Figure S9). Venom 302-like peptides have been identified in *Z. natalensis* [49] and the proteomes of *N. nomurai* [38] and the cubozoan *C. fleckeri* [22]

Auxiliary proteins can also include various proteases that may facilitate diffusion of neurotoxins by breaking down the extracellular matrix in prey, and display cytolytic, gelatinolytic, caseinolytic, and fibrinolytic functions in cnidarians [119]. The most diverse auxiliary proteins in the four cerianthid transcriptomes match to astacin-like metalloproteases (M12A) with a total of 52 sequences between the four cerianthids. This includes transcripts with a close match to nematocyst expressed protein 6 (NEP-6), an astacin family metalloprotease previously reported from the starlet sea anemone *Nematostella vectensis* [120].

Additional metalloproteases, including neprilysin-like toxins (peptidase_M13_N domain), also found in the venom of *Cyanea capillata* [41], and glutaminyl-peptide cyclotransferases (peptidase_M28 domain) were also expressed within each species. Metalloprotease M12B containing domain proteases (zinc metalloproteinase-disintegrin and coagulation factor X-activating enzyme heavy chain) are also found in all four cerianthid species (13 total), but are categorized as hemostatic and hemorrhagic toxins (Section 2.2.2.), since, in snake venoms, these toxins disrupt capillary activity [120]. M12B metalloproteases have also been found in the venoms of *N. nomurai* [38] and the hydrozoan *Olindias sambaquiensis* [122].

## 3. Discussion

In this study we assembled de-novo transcriptomes of four members of Ceriantharia: *C. brasiliensis, I. nocturnus, P. borealis*, and *P. maua*, with BUSCO scores between 88.1-97.9% completeness (Table 1). From these transcriptomes, we identified a total of 525 venom-like genes between all four species using our customized bioinformatic pipeline, which are sorted into 135 clusters (124 orthologous clusters and 12 single-copy gene clusters) (Supplementary Figure S11). The venom-like gene profiles of the four cerianthids are similar in composition and generalized biological function, though the annotated number of toxin-like genes within each species is highly variable (69-182). Our four cerianthid toxin profiles are similar to previous transcriptome-based venom profiles for cnidarians, including the prevalence of ShK-domain containing toxins (e.g. [22,38,46,54]). While each species has a diversity of toxins within each of the seven functional categories, all toxin profiles were dominated by hemostatic and hemorrhagic toxins (30.4%-40.3%), mixed function enzymes (12.4-21.7%) and auxiliary venom proteins/peptides (14.5%-20.3%) followed by neurotoxins (7.2-17.4%), allergen and innate immunity toxins (2.2-12.9%), protease inhibitors (2.4-4.3%), and membrane-active toxins (0-5.7%). It should be noted that many of these toxins may have alternative or additional molecular functions, and the presented categorization only represents broad patterns based on previous studies on animal venoms. There was also a significant proportion of “unknown” toxins from each species within each transcriptome assembly (Table 2, Figure 2). Given that this is the first survey of putative toxins in this subclass within an already understudied group, it is unclear if these unknowns are potential novel venom-like transcripts or potential artifacts of assembly and annotation.

**Table 2.**
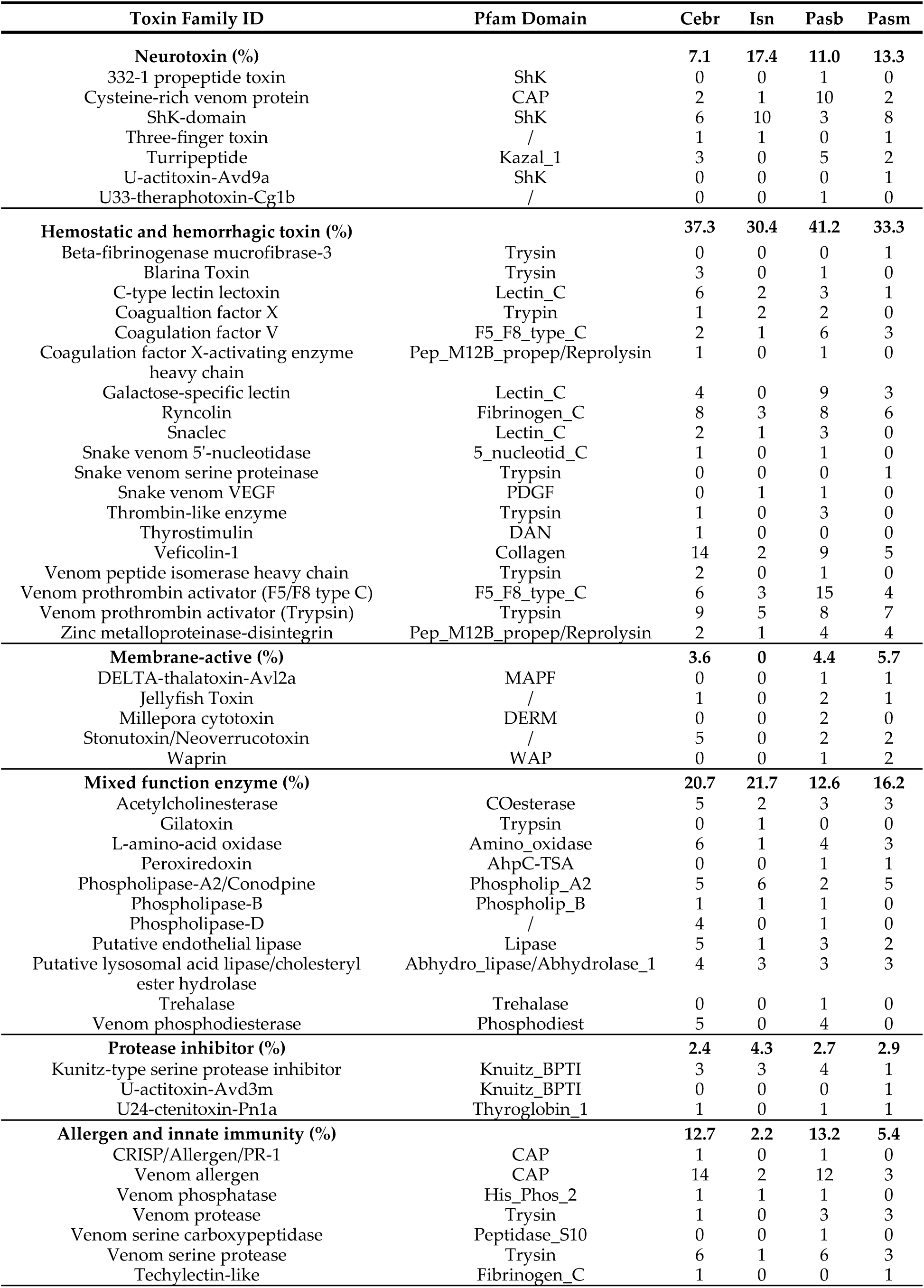
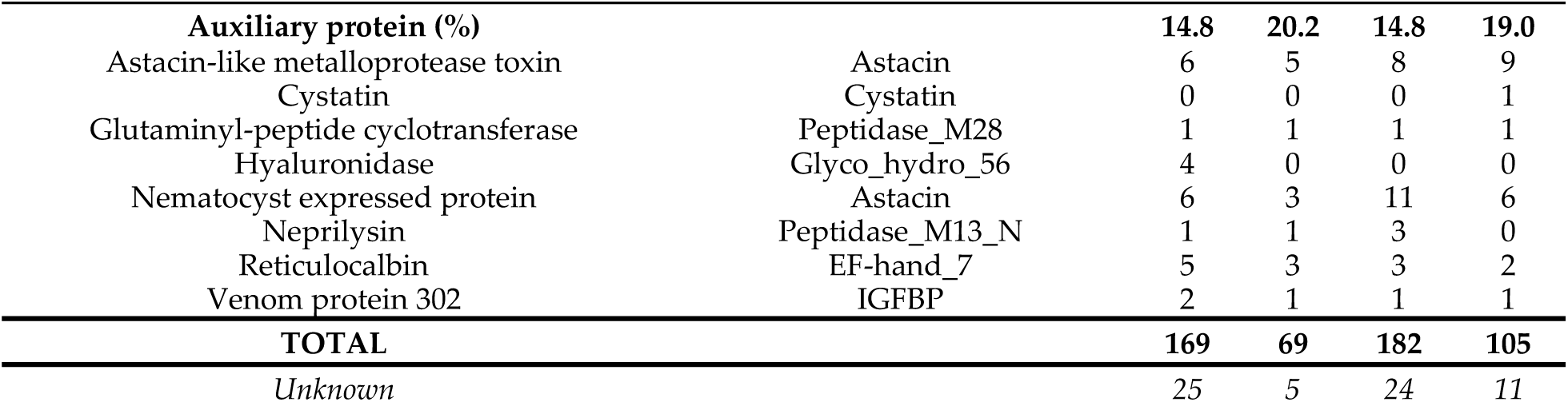
Toxin families identified for each cerianthid species.

Some of the most common families we identified are common in anthozoan venoms, including PLA2, metalloproteases, serine proteases, and kunitz-domain protease inhibitors [11,43,51]. Several of the less common venom-gene families identified in cerianthids have also been identified in the transcriptomes of colonial zoanthids [47-49], another understudied group of anthozoans, including turripeptides, three-finger toxins, and venom protein 302 toxins (Figure 4; Supplementary Figure S4,S9), as well as snake venom VEGF toxins. However, the phylogenetic evidence for the majority of these candidate toxins is weak due to clustering with non-venomous taxa and/or low bootstrap scores. As mentioned above, several of these toxin groups have been identified in other cnidarian groups, including turripeptides [22,82,83] and venom protein 302 [22,38]. It is unclear if the similarities of these less common toxin families between zoanthid and cerianthid toxins are due to shared biology/evolutionary history or an artifact of the relatively limited dataset for cnidarians.

While membrane-active or pore-forming toxins are common in most cnidarian venoms [123], we had not expected to capture putative toxins in the jellyfish toxin family (also called CaTX/CrTX toxin family) in three of the four cerianthids species, given that these toxins are primarily found in medically relevant cubozoan venoms (Figure 6). In an ecological context, these highly potent toxins likely allow box jellyfish to capture fish [124,125]; while the diet of cerianthids remains fairly ambiguous, it is unlikely they capture fish as prey. Toxins from this family have previously been identified in other anthozoan species through genomic and transcriptomic studies [40,51,126], but to the best of our knowledge, these toxins have never been detected through proteomic methods in anthozoans [40]. Given that these toxins are present in multiple cerianthids (including two paralogs within *P. borealis*), these toxins are good candidates for proteomic analysis and potentially functional characterization.

Because cerianthids group within the class Anthozoa, it is interesting that several toxins groups commonly reported in anthozoans were absent from all four cerianthid species. For example, we expected to find a diverse set of low molecular weight neurotoxins, such as sea anemone sodium (Na+) channel toxins, potassium (K+) channel toxins, small cysteine-rich peptides (SCRiPs), sodium-selection acid-sensing ion channel (ASICs) inhibitors, and nonselective cation channel (TRPV1) inhibitors [11,25,30,127]. However, the four cerianthids transcriptomes contained relatively low numbers of neurotoxins in general, and only a single transcript from *P. maua* closely matched a sea anemone type I K+ channel toxin (Table 1). Additionally, actinoporin-like sequences are often found in sea anemones and other organisms [93,123], but only two actinoporin-like sequences were found in *P. borealis* and *P. maua*, despite often being found in sea anemones. We also found no evidence of small cysteine-rich peptides (SCRiPS), neurotoxins with eight conserved cysteine residues that cause paralysis in zebrafish (*Danio rerio*) [128], which were initially reported in the corals *Orbicella faveolata* (as *Montastraea faveolata*), *Montipora capitata*, and *Acropora millepora* [129]. The vast majority of candidate toxins containing ShK domains did not have a close match to any toxin in the Tox-Prot database, but in 22 sequences we could confidently determine the six cysteine residue patterns characteristic of ShK domains (Supplementary Figure S3). The exponential increase in ShK domain peptides found in anthozoans prompted a recent sequence-function study of the superfamily [130], and cerianthid ShK-domain toxins may represent additional structural scaffolds with novel function for further study.

In general, our findings contrast with the previously observed pattern that anthozoan venoms are typically neurotoxin-rich while medusozoan venoms are dominantly enzymatic. The venoms of anthozoans and medusozoans have been broadly reported to be distinct, with hydrozoans, scyphozoan, and cubozoan venoms being dominated by larger cytolytic proteins and anthozoans by low molecular weight neuropeptides [26,40,83]. However, this pattern is based on highly biased taxonomic data, as mentioned above [27]. Even though a greater diversity of enzymatic-like genes is present within the four cerianthid transcriptomes, it is possible the level of protein expression could shift towards a smaller subset of toxins dominating the venom composition, and therefore overall venom function. For example, it has been shown in *S. haddoni* that even when more enzymatic toxin-like sequences are present in the transcriptome, the expression of neurotoxins is greater overall in milked venom (i.e. the proteomic level) [46]. Thus, future quantitative gene expression and proteomic studies are needed to provide a more holistic understanding of both single toxin and whole venom function in these species.

Because the phylogenetic placement of Subclass Ceriantharia remains unclear, it is difficult to interpret the evolutionary context of their venom profile within Anthozoa. For instance, if Ceriantharia is sister to the Hexacorallia, that suggests that the expansion of neuropeptide toxins occurred after the divergence of Ceriantharia, possibly through extensive gene duplications [52,126,131]. Neurotoxins in sea anemones are important because they are sessile animals, and may be critical to deterring predators [132]. Because cerianthids can fully contract into their tubes, they have a distinct means of protecting themselves from predators in contrast to sea anemones which cannot fully retract their bodies, which may ease the selective pressure to diversify or maintain defensive toxins. If Ceriantharia is instead sister to Hexacorallia + Octcorallia, families such as the jellyfish toxins may have been present in the last common ancestor and subsequently lost in the other anthozoan lineages. Additionally, as noted above, cerianthids often have a long-lasting pelagic larval stage. There is a general consensus that the composition and function of toxins reflects the ecological utility of that venom [133], thus, the increased time in the pelagic environment in the larval stage likely exposes cerianthids to different sets of potential predators and prey, resulting in different selection pressures driving venom composition and function. We can only speculate on the role of these various venom components and overall venom function in the ecological interactions of these animals until additional molecular studies are completed [27,134].

One interesting outcome is the difference in the number of venom-like putative protein coding transcripts found in *I. nocturnus* compared to the other three species (69 compared to 169, 182, 105). As this species is the only representative of the family Arachnactidae, this may be evidence of evolutionary difference compared to the family Cerianthidae, which is corroborated by morphology and traditionally accepted [73]. At the ecological level, the species *I. nocturnus*, as its name indicates, is nocturnal and thus increases its activity at night. This may indicate different needs in relation to predation and prey capture compared to species active during the day. For instance, species of the family Arachnactidae show considerable concentrations of green fluorescent protein [135], which can be an important mechanism of prey capture at night [136]. This may relax the selective pressures, or potentially the available metabolic energy, to sustain a large, complex toxin arsenal, and therefore result in the lower number of venom-like genes identified in our study.

While our findings suggest several interesting patterns about presence and absence of certain cerianthid venom components, there are some limitations to exploring the venom profiles of understudied species. Previous studies have shown that cnidarian transcriptomes often yield a larger diversity of putative toxin sequences than a combined transcriptome-proteome approach (e.g. [46,53,54,126]). This difference may be reflective of the state of the animal when collected; animals that have recently fired their stinging cells will likely express more venom-like genes as venom is being synthesized for developing nematocysts [46]. Consequently, animals that have not discharged their stinging cells recently may have a lower than expected expression of toxin-like sequences. There are also often issues using de-novo assemblies for venom gene discovery, including high false discovery rate or inability to annotate novel venom genes [137,138]. For instance, even though no membrane-active toxins were detected in *I. nocturnus*, it is unlikely that there are truly no toxins with this function, especially given their ubiquity in cnidarians [139]. Our study also focused on candidate transcripts that contained full ORFs (stop and start codon), which likely decreased the diversity of toxin-like gene candidates. The set of venom-like genes we present here are viewed as an initial step into exploring the diversity of the toxin peptides and proteins within a poorly studied cnidarian group.

We present the first sequence-based analysis of venom-like genes within the Subclass Ceriantharia. The four species of cerianthids expressed over 500 novel toxin-like genes that are functionally and structurally diverse. While the overall functional profiles are similar to other transcriptomic studies of cnidarians, many common anthozoan toxin families are not present in our study. This could have notable implications both for the evolution of venom genes with Anthozoan as well as ecological utility of candidate toxins within this specific anthozoan lineage. Furthermore, the addition set of ShK-domain, as well as kunitz-domain containing toxins, shows that cerinathid toxins provide potential candidates for therapeutic study. We hope that these new data will be utilized to further explore the diversity and function of these venom proteins and peptides.

## 4. Materials and Methods

### 4.1. Tissue collection, RNA extraction, next-gen sequencing, and transcriptome assembly

Four species were used in the current study. The species C. brasiliensis and I. nocturnus were obtained in São Sebastião, São Paulo, Brazil while SCUBA Diving. The P. borealis specimen was purchased through Gulf Of Maine inc. (Pembroke, ME). The P. cf. maua specimen was purchased from an aquarium supplier and currently on exhibit at Discovery Place Science (Charlotte, NC). For each species, several (10+) tentacles were collected from each organism after acclimating them to aquariums for 48 hours or longer. Tissues were flash frozen in liquid nitrogen or stored in RNA later in -80°C. Total RNA was extracted using the RNAqueous Total RNA Isolation Kit from Thermo Fisher Scientific (Massachusetts, USA). RNA was assessed using a NanoDrop 2000 spectrophotometer (Thermo Fisher). High throughput Sequencing was done on an Illumina HiSeq at the DHMRI (Kannapolis, NC). Total RNA was quantitated using the Quant-iT RiboGreen RNA Assay Kit (Thermo Fisher) and RNA integrity assessed using the Agilent Bioanalyzer. RNA sequencing libraries were generated using the Illumina TruSeq RNA Library Prep RNA Kit following the manufacturer’s protocol and quantitated using qPCR and fragments visualized using an Agilent Bioanalyzer. Libraries were combined in equimolar amounts onto one flow cell for a 125 bp paired end sequencing run on the Illumina HiSeq2500. Overall quality of the sequencing run evaluated using FastQC [140]. Transcriptome assembly was done using the de novo assembly program Trinity v2.2 [74]. Transcriptome completeness was determined using the program BUSCO v3 [141].

### 4.2. Bioinformatic analysis and venom annotation

For the custom annotation pipeline, protein-coding regions were predicted from assembled transcriptomes using TransDecoder v5.5.0, minimum set to 50 (https://transdecoder.github.io) [142]. Using blastp from NCBI BLAST+ v.2.8.1 [143,144] with an e-value cutoff of 0.001, all transcripts were searched against 1) proteins and toxins from the Tox-prot animal venom annotation database ([57], downloaded March 2019), and 2) all cnidarian toxins and proteins from the Protein database on NCBI (“Cnidaria AND ((Toxin) OR (Venom)),” downloaded March 2019). Additionally, predicated protein-coding regions were searched using hmmsearch with an e-value cutoff of 0.001 from HMMER 3.1b2 [145,146] against hidden markov model (HMM) profiles from alignments of 20 venom protein classes. HMM were modified from those used in a transcriptomic study on the venom of bloodworms [81] by supplementing several cnidarian specific toxins within respective venom protein families. Additionally, four cnidarian-specific pore-forming venom families were added based on annotations from VenomZone (venomzone.expasy.org, accessed March 2018): Actinoporin sea anemone subfamily, jellyfish toxin family, cnidaria small cysteine-rich protein (SCRiP) family and MACPF-domain toxins. The results from all three searches were combined and all complete coding sequences used for downstream analysis. Venoms are secreted proteins and peptides, thus signal peptides were predicted using the SignalP v5.0 server (https://services.healthtech.dtu.dk/service.php?SignalP-5.0) [147]. Redundant sequences were clustered using CD-HIT v.4.6.8 with a cutoff of 0.95 [148,149]. A reciprocal search using blastp was used with an e-value cutoff of 1e-5 against Tox-Prot animal venom database and the NCBI non-redundant protein sequences (nr) database (downloaded March 2019), as well as a hmmsearch search with an evalue cutoff of 1e-5 against Pfam (downloaded March 2019) [77].

The results were manually curated to confirm that BLASTp annotations from ToxProt matched the detected venom domain from Pfam [76,77]. In addition, several toxins were not identified from ToxProt that were from NCBI database (e.g. three-finger toxin W-IV-like (NCBI Reference Sequence: XP_015758456.1), 332-1 secreted propeptoide (GenBank: AKU77030.1). Candidates were considered “unknown” and not used for further analysis if there was no match to a protein from Tox-Prot, the best match from NCBI was an uncharacterized or predicated protein, and no toxin domain was detected. The final list of candidate toxins was classified into protein families, molecular function (based on annotation from UniProtKB/Swiss-Prot) [150], and putative biological function. The results were visualized using the PieDonut via the webr package v.0.1.2 (https://cardiomoon.github.io/webr/) in R v3.6.2 [151] within Rstudio v1.0.153 [152] and final figures constructed in Inkscape v1.0beta2 (inkscape.org).

### 4.3. Phylogenetic analysis of select gene families

For select toxin families, gene trees were constructed using a representative set of venomous and non-venomous proteins for each protein family, modified from phylogenetic analyses in von [80] and [47]. Candidate cerianthids toxins and were aligned using the L-INS-I algorithm in MAFFT v7.312 [153]. Maximum likelihood phylogenies were constructed using RAxML v8.2.12 [154] under the PROTGAMMA + WAG model and branch support calculated using 500 rapid bootstrap replicates (-x). Trees were visualized using FigTree v1.4.4 (https://github.com/rambaut/figtree) and final figures constructed in Inkscape v1.0beta2 (inkscape.org).

### 4.4. Availability of supporting data

Raw reads used to construct the transcriptomes used in this analysis have been deposited under the SRA bioproject PRJNA633022, specifically SRR11802642 (*C. brasiliensis*), SRR11802641 (*I. nocturnus*), SRR11802643 (*P. borealis*), and SRR11802640 (*P. maua*) accessions.

## Supplementary Materials

Figure S1: Bioinformatic pipeline for the annotation of venom-like genes for four cerianthid transcriptomes, Figure S2-S10: Phylogenetic relationships between several toxin gene families and putative cerianthid sequences, Figure S11: Orthologous gene clusters of the putative venom-like genes for all four cerianthids, Table S1: Annotation table for putative venom-like genes for four cerianthid species (Excel).

## Author Contributions

SNS, JM and AMR obtained samples, JM and SNS extracted RNA, JM assembled transcriptomes, AMLK performed the analysis and annotation of toxins and wrote the initial draft. AMLK, SNS, JM and AMR contributed to the manuscript.

## Acknowledgments

We are grateful to Elliot Provance, Drs Marymegan Daly, André C. Morandini, Paulyn Cartwright, Marcelo V. Kitahara, Joe Ryan, and Melissa Debiasse for assisting with obtaining materials and/or discussions about the study.

