## Supplemental Figures for "Transcriptomic analysis of four cerianthid (Cnidaria, Ceriantharia) venoms"

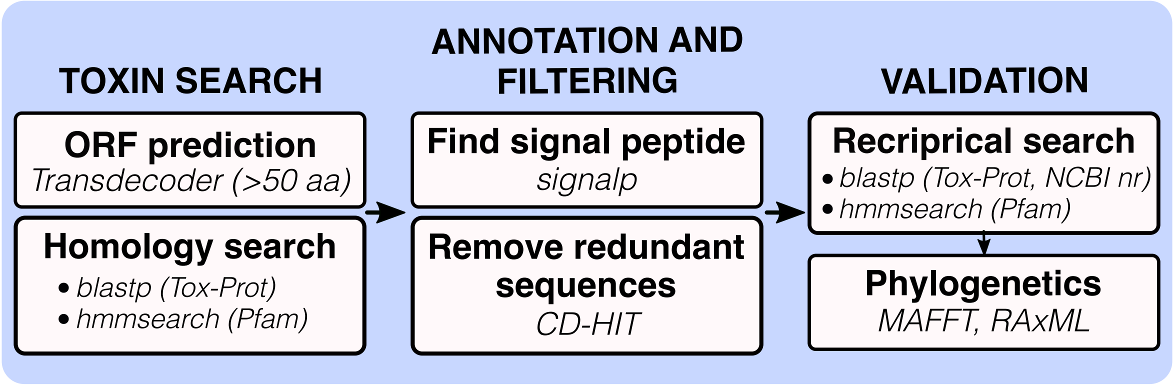


**Figure S1.** Bioinformatic pipeline for the annotation of venom-like genes for four cerianthid transcriptomes.

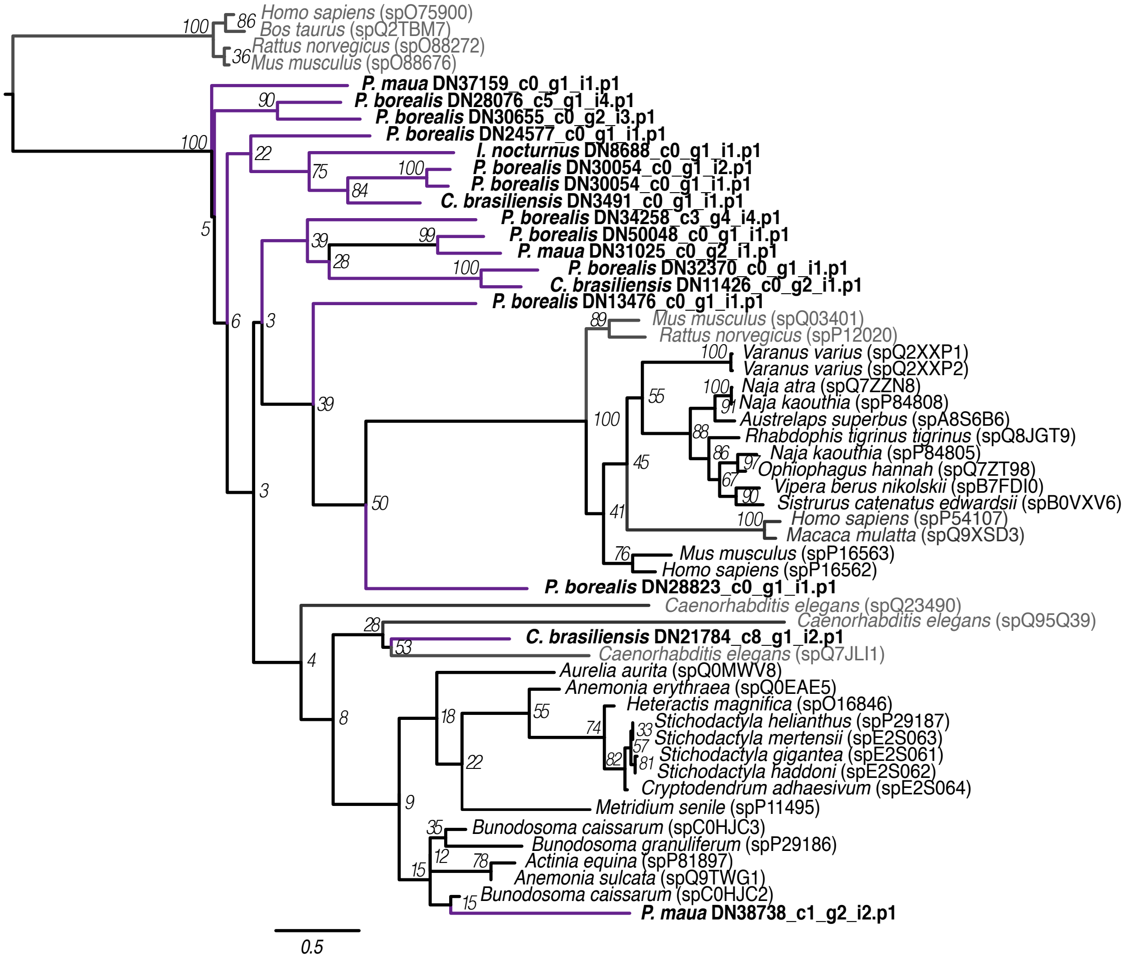


**Figure S2.** Phylogenetic tree of cystine-rich protein family sequences. Phylogeny was constructed using RAxML with the PROTGAMMAWAG option. Bootstrap support based on 500 rapid bootstrap replicates, and all support values are shown. Sequence names in bold with purple branches are those from cerianthids. Sequences in gray are non-venomous representatives. Phylogeny modified from [81].

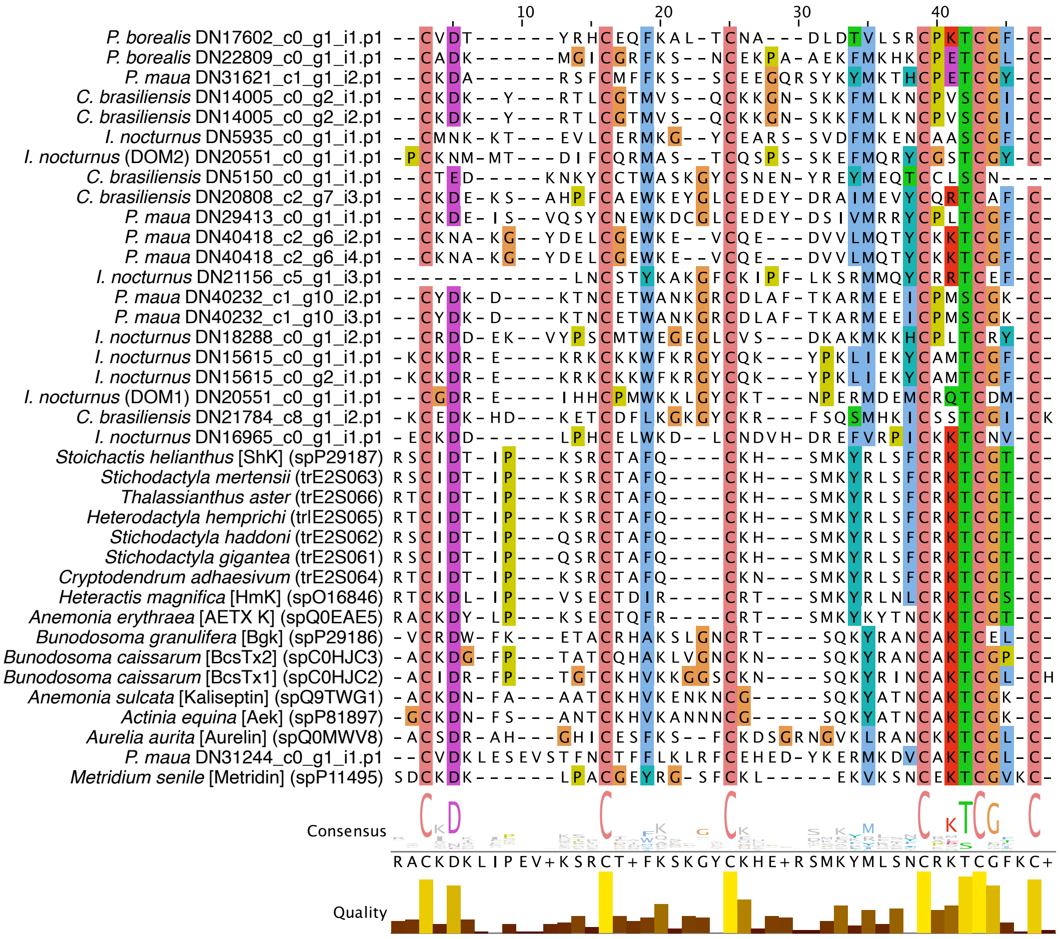

**Figure S3.** Multiple sequence alignment of ShK domains in cerianthid venom-like genes and other toxin representatives. Sequences alignment constructed using the L-INS-I algorithm in MAFFT [153] and viewed in a Clustal color scheme using Jalview v2.11.1 [155]. Six conserved cystine residues characteristic of ShK domains are should across all taxa. Various SHK toxin domains modified from [81].

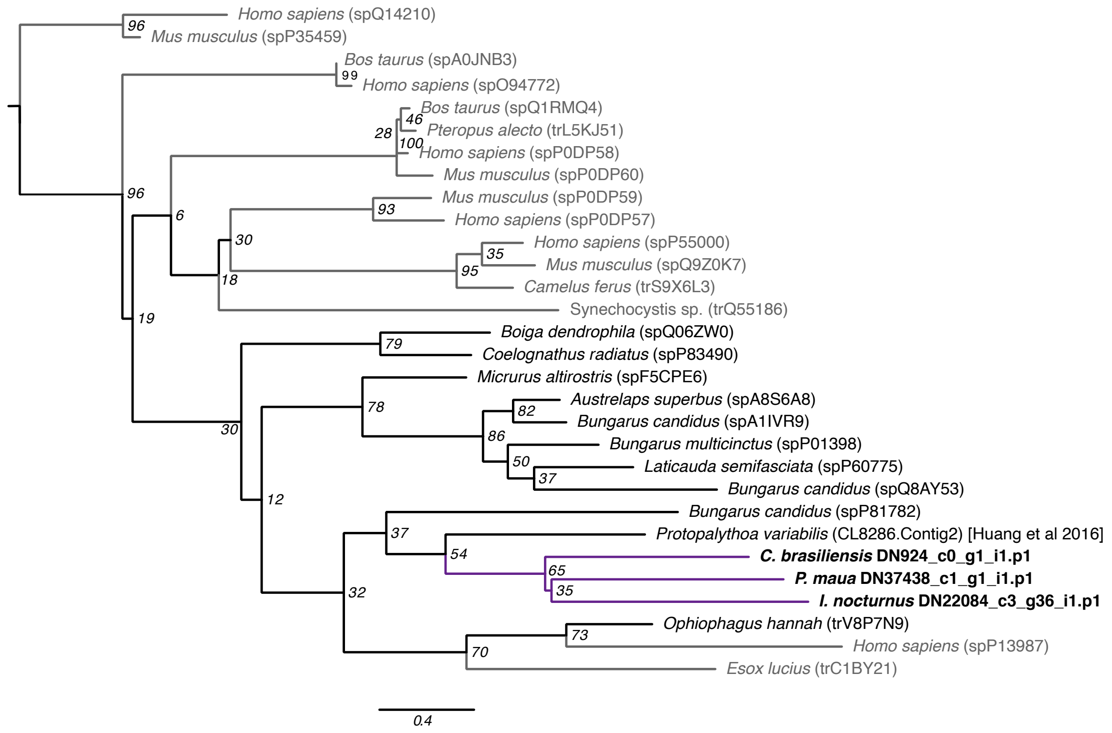


**Figure S4.** Phylogenetic tree of three-finger toxin family sequences. Phylogeny was constructed using RAxML with the PROTGAMMAWAG option. Bootstrap support based on 500 rapid bootstrap replicates, and all support values are shown. Sequence names in bold with purple branches are those from cerianthids. Sequences in gray are non-venomous representatives. Phylogeny modified from [47].

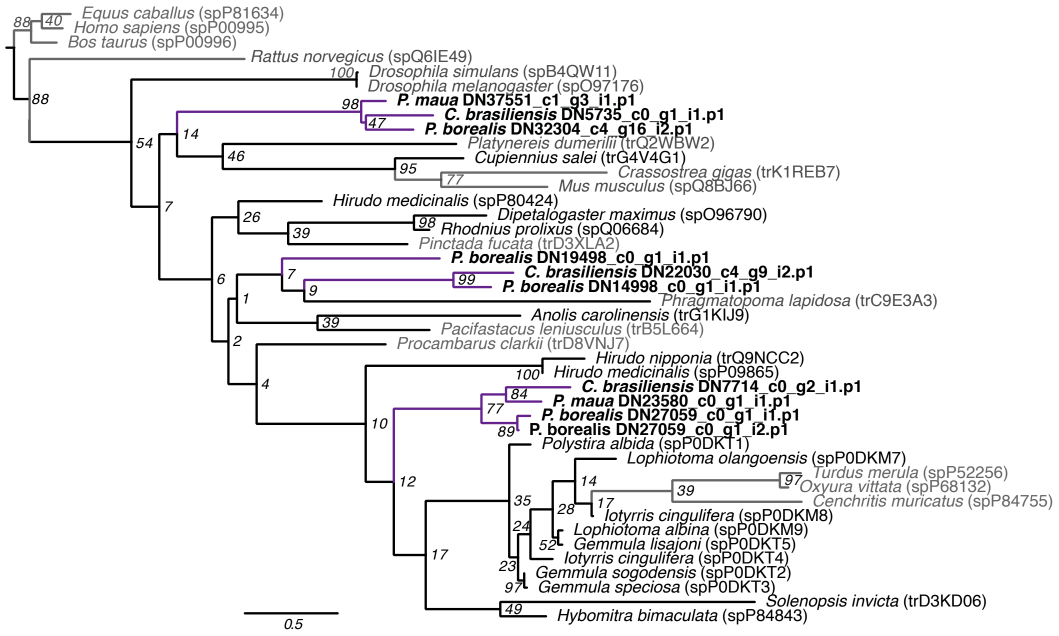


**Figure S5.** Phylogenetic tree of kazal family sequences. Phylogeny was constructed using RAxML with the PROTGAMMAWAG option. Bootstrap support based on 500 rapid bootstrap replicates, and all support values are shown. Sequence names in bold with purple branches are those from cerianthids. Sequences in gray are non-venomous representatives. Phylogeny modified from [81].

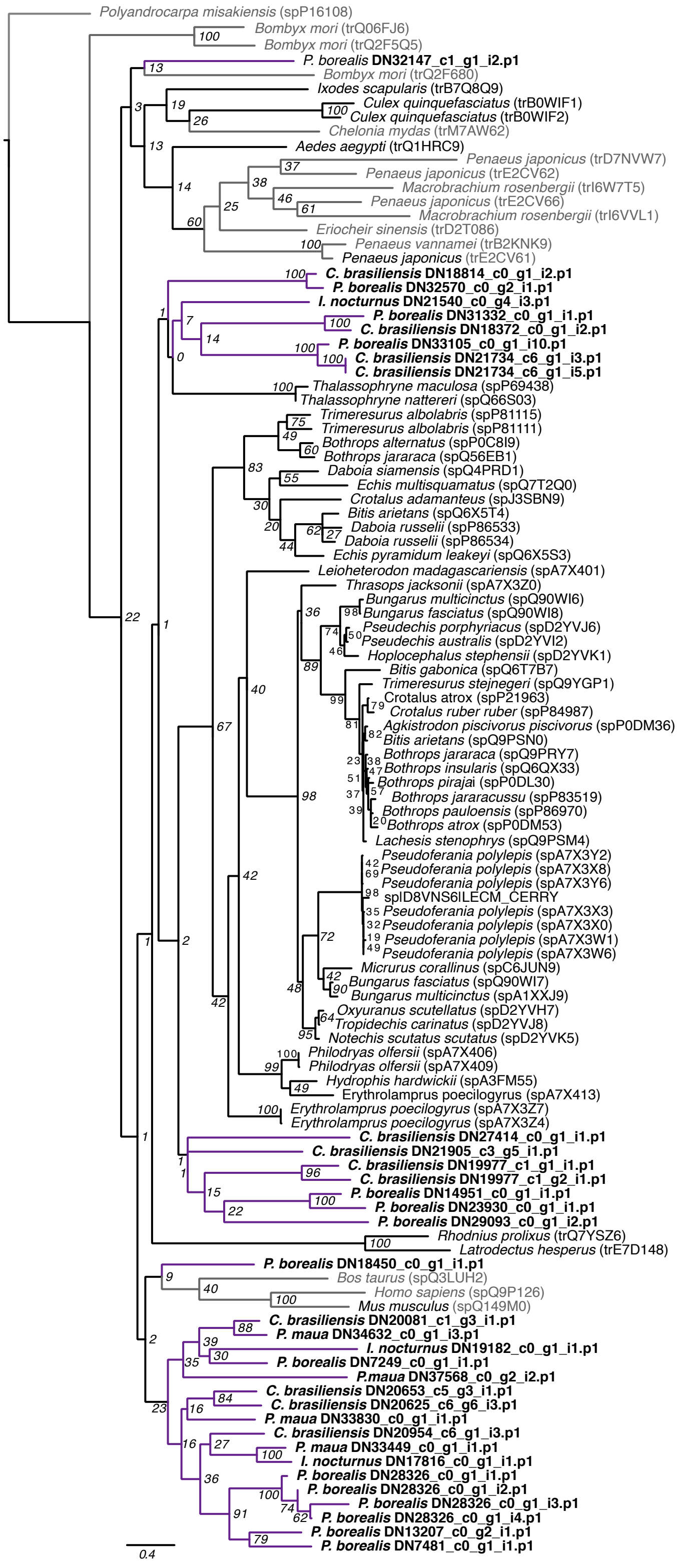


**Figure S6.** Phylogenetic tree of C-type lectin family sequences. Phylogeny was constructed using RAxML with the PROTGAMMAWAG option. Bootstrap support based on 500 rapid bootstrap replicates, and all support values are shown. Sequence names in bold with purple branches are those from cerianthids. Sequences in gray are non-venomous representatives. Phylogeny modified from [81].

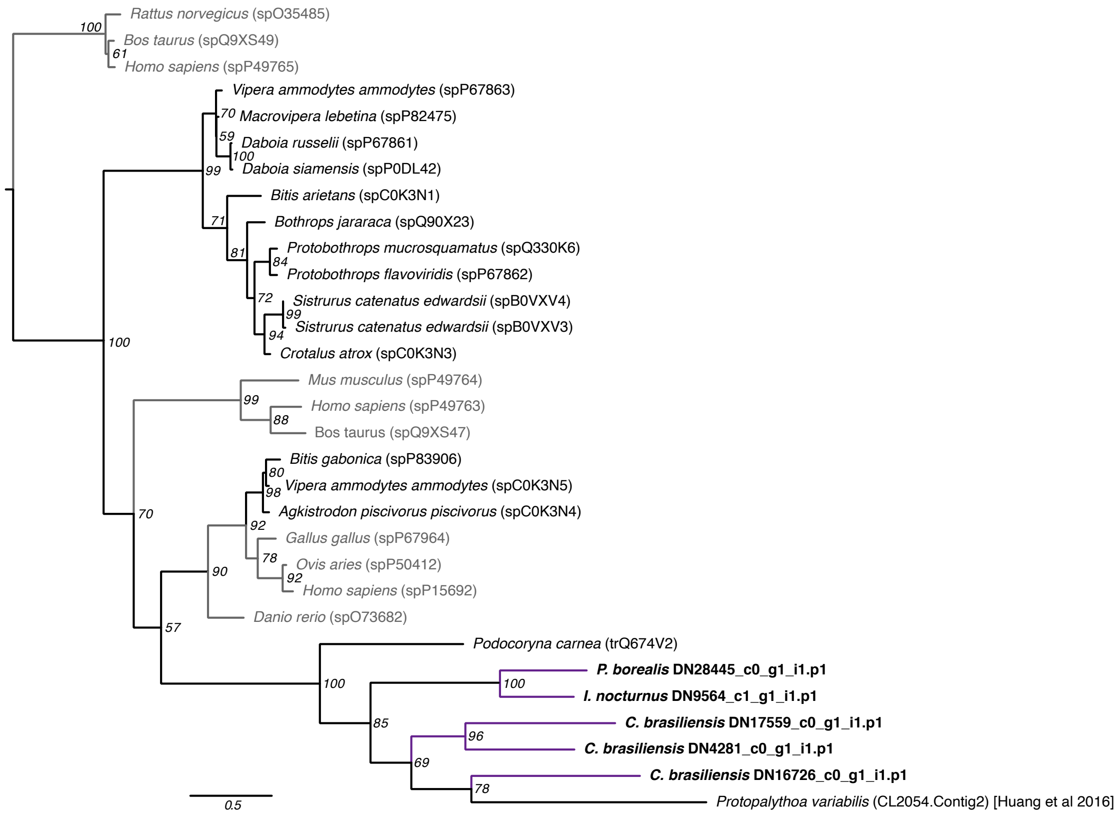


**Figure S7.** Phylogenetic tree of venom vascular endothelial growth factors (venom VEGF) family sequences. Phylogeny was constructed using RAxML with the PROTGAMMAWAG option. Bootstrap support based on 500 rapid bootstrap replicates, and all support values are shown. Sequence names in bold with purple branches are those from cerianthids. Sequences in gray are non-venomous representatives. Phylogeny modified from [47].

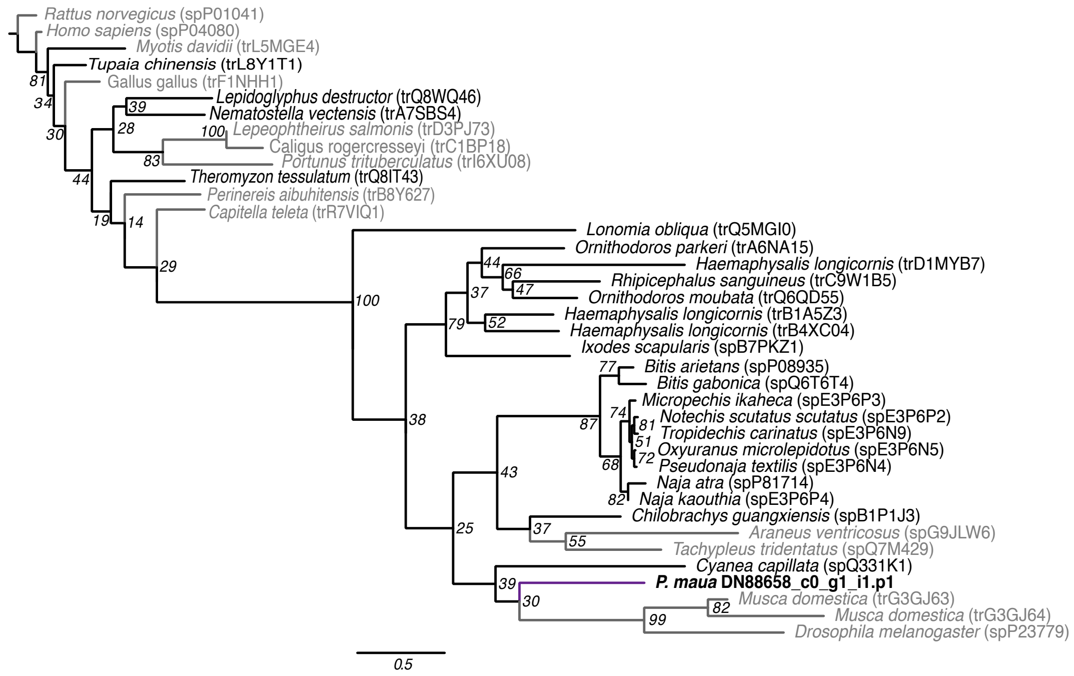


**Figure S8.** Phylogenetic tree of cystatin family sequences. Phylogeny was constructed using RAxML with the PROTGAMMAWAG option. Bootstrap support based on 500 rapid bootstrap replicates, and all support values are shown. Sequence names in bold with purple branches are those from cerianthids. Sequences in gray are non-venomous representatives. Phylogeny modified from [81].

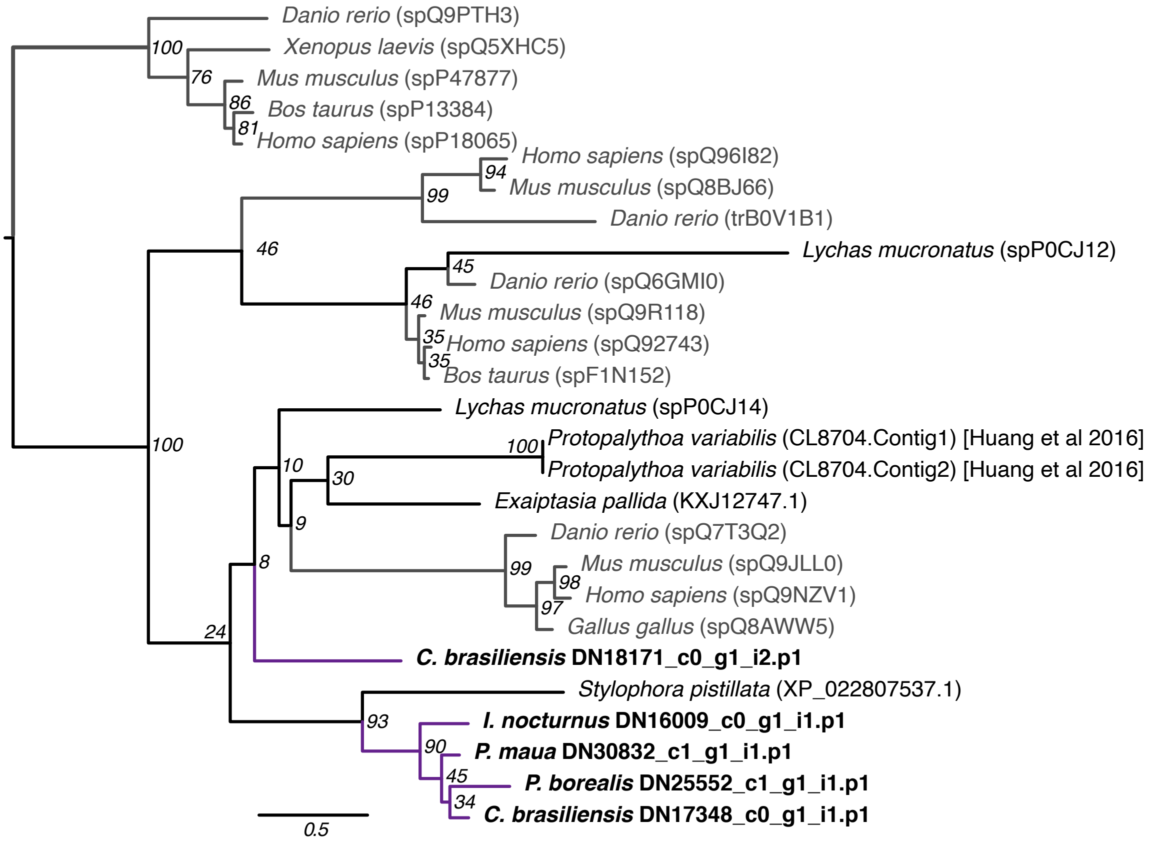


**Figure S9.** Phylogenetic tree of venom protein 302 family sequences. Phylogeny was constructed using RAxML with the PROTGAMMAWAG option. Bootstrap support based on 500 rapid bootstrap replicates, and all support values are shown. Sequence names in bold with purple branches are those from cerianthids. Sequences in gray are non-venomous representatives. Phylogeny modified from [47].

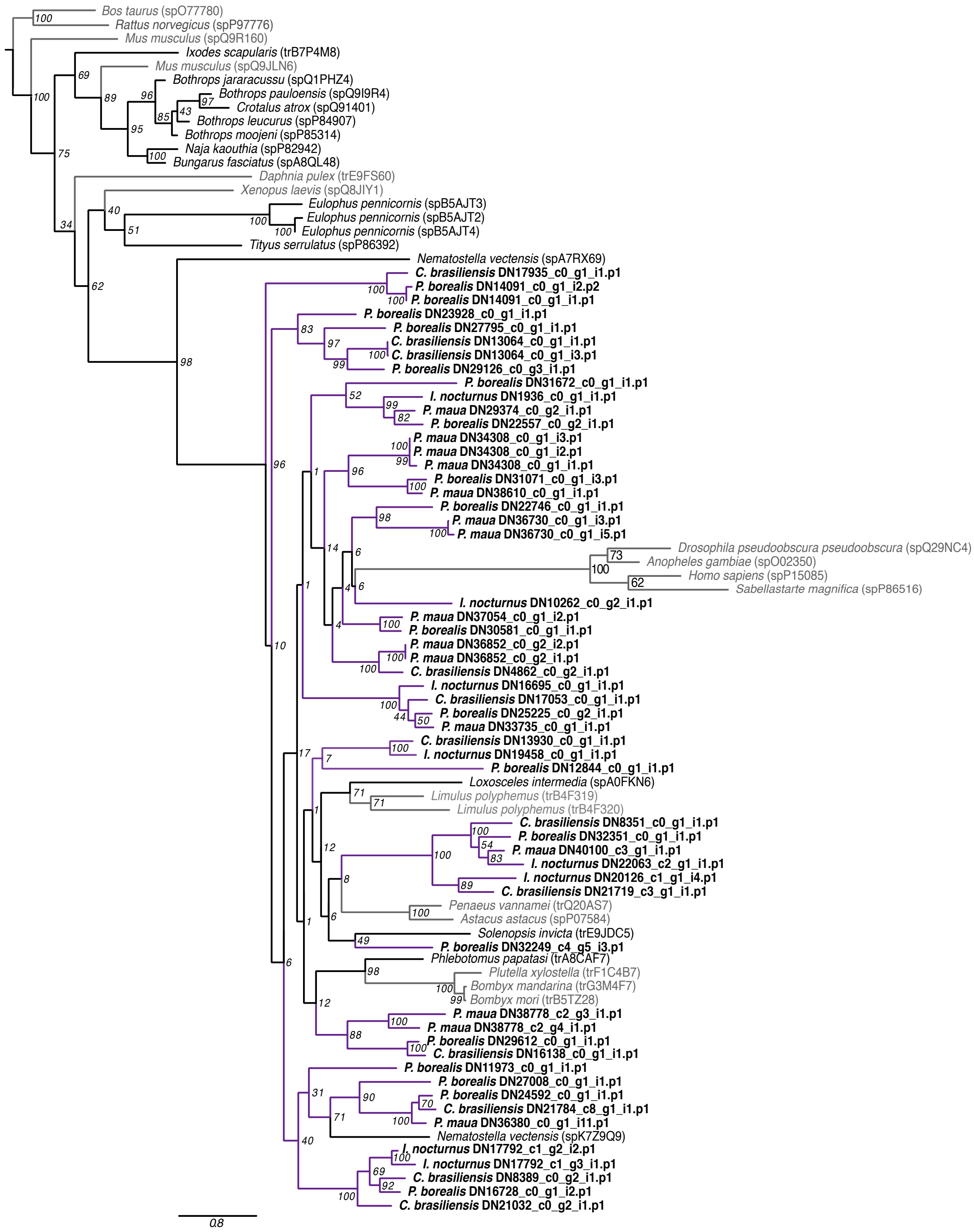


**Figure S10.** Phylogenetic tree of metalloprotease M12A family sequences. Phylogeny was constructed using RAxML with the PROTGAMMAWAG option. Bootstrap support based on 500 rapid bootstrap replicates, and all support values are shown. Sequence names in bold with purple branches are those from cerianthids. Sequences in gray are non-venomous representatives. Phylogeny modified from [81].

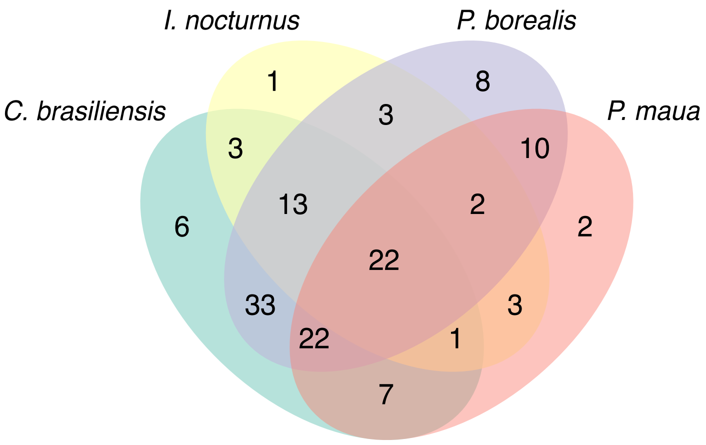


| Species | # of proteins, (in cluster) | # of proteins, (singletons) |
|---|---|---|
| *C. brasiliensis* | 107 | 27 |
| *I. nocturnus* | 48 | 13 |
| *P. borealis* | 113 | 43 |
| *P. maua.* | 69 | 18 |

**Figure S11.** Orthologous gene clusters of the putative venom-like genes for all four cerianthids. Venn diagram shows overlap of gene clusters for all four species. Table shows the number of venom-like genes within each species that fall within a cluster or are singletons, which as genes that cannot be placed into a cluster (unique to each species). Gene clusters were analyzed using the OrthoVenn2 server under default setting except evalue changed to 1e-5 (https://orthovenn2.bioinfotoolkits.net/) [156].
